# C-terminal truncation of Pik3r1 in mice models human lipodystrophic insulin resistance uncoupled from dyslipidemia

**DOI:** 10.1101/224485

**Authors:** Albert Kwok, Ilona Zvetkova, Sam Virtue, Isabel Huang-Doran, Patsy Tomlinson, David A. Bulger, Daniel Hart, Rachel Knox, Stephen Franks, Peter Voshol, Antonio Vidal-Puig, Amanda N. Sferruzzi-Perri, Jørgen Jensen, Stephen O’Rahilly, Robert K Semple

**Affiliations:** The University of Cambridge Metabolic Research Laboratories, Wellcome Trust-MRC Institute of Metabolic Science, Cambridge, UK; The National Institute for Health Research Cambridge Biomedical Research Centre, Cambridge, UK; MRC Metabolic Diseases Unit, Wellcome Trust-MRC Institute of Metabolic Science, Cambridge, UK; Institute of Reproductive & Developmental Biology, Department of Surgery & Cancer, Imperial College London, London, UK; Louis Bolk Institute, Kosterijland 3-5, NL-3981AJ, Bunnik, The Netherlands; Centre for Trophoblast Research, Department of Physiology, Development and Neuroscience, Downing Street, University of Cambridge, Cambridge, UK; Department of Physical Performance, Norwegian School of Sport Sciences, P.O.Box 4014, Ulleval Stadion, 0806 Oslo, Norway

## Abstract

Heterodimeric class IA phosphatidylinositol-3-kinases (PI3K) transduce signals from many receptor tyrosine kinases including the insulin receptor. PI3K recruitment to phosphotyrosines is mediated by *Pik3r1* gene products including the most intensely studied PI3K regulatory subunit, p85α, which also binds and regulates the PIP3 phosphatase *Pten,* and the lipogenic transcription factor *Xbp1.* Mutations in human *PIK3R1* cause SHORT syndrome, featuring lipodystrophy and severe insulin resistance which, uniquely, are uncoupled from fatty liver and dyslipidemia. We describe a novel mouse model of SHORT syndrome made by knock in of the *Pik3r1* Y657X mutation. Homozygous embryos die at E11.5, while heterozygous mice exhibit pre-and postnatal growth impairment with diminished placental vascularity. Adipose tissue accretion on high fat feeding was reduced, however adipocyte size was unchanged and preadipocyte differentiation *ex vivo* unimpaired. Despite severe insulin resistance, heterozygous mice were hypolipidemic, and plasma adiponectin, liver weight, cholesterol, glycogen and triglyceride content were unchanged. Mild downregulation of lipogenic *Srebp1, Srebp2* and *Chrebp* transcriptional activity but no suppression of *Xbp1* target genes was seen after fasting. These findings give new insights into the developmental role of *Pik3r1,* and establish a model of lipodystrophic insulin resistance dissociated from dyslipidemia as seen in SHORT syndrome.

## INTRODUCTION

Class 1A phosphatidylinositol-3-kinases (PI3K) phosphorylate phosphatidylinositol-4,5-bisphosphate to produce phosphatidylinositol-3,4,5-trisphosphate, or PIP3. They are composed of a heterodimer of a p110 catalytic subunit (either p110α, β or δ, encoded in humans by *PIK3CA, PIK3CB* and *PIK3CD* respectively) tightly bound to a regulatory subunit, three of which - p85α, p55α or p50α - are encoded by the *PIK3R1* gene. Each of these *PIK3R1* gene products is able to bind any of the three catalytic subunits with no discernible selectivity, stabilising them and mediating recruitment of PI3K holoenzymes to activated receptor tyrosine kinases or their substrates through binding of one or both SH2 domains to phosphotyrosines within YMXM motifs (1, 2).

The critical importance of Class 1A PI3K in human physiology is manifest in the convincing association of activating mutations in catalytic subunits with disease: activating mutations in *PIK3CA,* encoding p110α, occur at high frequency both in cancers (3), and in a wide range of asymmetric forms of overgrowth, where mutations occur postzygotically and are in a mosaic distribution (4). Genetic and pharmacological studies have moreover established that p110a is critical in transducing the metabolic actions of insulin (5, 6), however it is also coupled to a wide range of other receptor tyrosine kinases. p110δ is predominantly important in lymphocytes, and activating mutations cause autosomal dominant immunodeficiency (7), while the role of the ubiquitous p110β has been less precisely defined (8).

Genetic efforts to probe the role of PIK3R1-encoded regulatory subunits have yielded a more complex picture. Selective deficiency of p85α in mice or humans leads to immunodeficiency (9, 10) and enhanced insulin sensitivity (11), with the latter also seen in mice with heterozygous p85α deletion (12) or deletion of both p50α and p55α subunits (13). Deficiency of all three gene products leads to perinatal mortality in mice with liver necrosis (14), and when the p85β regulatory subunit, encoded by *Pik3r2*, is also knocked out, embryos die around E12.5 with evidence of failure of turning (15). Collectively these findings demonstrate that *Pik3r1* gene products and p85β have some redundant functions. The enhanced insulin sensitivity seen in p85α deficiency has been argued to be due to the presence of excess free p85α subunits in the wild-type state which can compete with PI3K heterodimers at phosphotyrosines (16), although there is some evidence to counter this notion, and other possibilities such as alterations in PIP3 phosphatase activity have been advanced (17).

*PIK3R1* gene products have also been suggested to have signaling roles beyond stabilising PI3K and conferring recruitability to phosphorylated YMXM motifs. Pertinent to metabolic homeostasis, for example, is the ability of p85α in mice to bind the transcription factor Xbp1 and to traffic it to the nucleus, where it regulates transcription of effector genes of the unfolded protein response (18, 19), as well as lipogenic genes (20).

In the face of this complexity, the discovery that missense or nonsense mutations in the C-terminal SH2 domain of *PIK3R1,* which affect all three protein products of the gene, produce SHORT syndrome (21–23), was of great interest. SHORT syndrome is named according to visible dysmorphic features (short stature, hyperextensibility of joints and/or hernia, ocular depression, Rieger anomaly of the iris, and teething delay), however it is the associated metabolic disorders that are potentially more informative about mechanisms of pandemic metabolic disease. Most patients with SHORT syndrome show insulin resistance that is often very severe, and partial lipodystrophy is also common (24). Highly unusually, lipodystrophy and severe insulin resistance are uncoupled from fatty liver and metabolic dyslipidaemia in this setting, for reasons that are not clear, and adiponectin concentrations in the plasma are preserved, unlike in pandemic insulin resistance (25). Affected women are also commonly anovulatory with severely elevated testosterone concentrations in the blood (25).

Recently, mice heterozygous for the commonest SHORT syndrome-associated *PIK3R1* mutation, R649W, were reported to exhibit features of SHORT syndrome including reduced linear growth, partial lipodystrophy, and systemic insulin resistance (26). We now report a novel murine model of SHORT syndrome created by knock in of a different pathogenic human allele, Y657X, that severely truncates the C terminal SH2 domain of all PIK3R1 gene products. We extend previous phenotyping to demonstrate the consequences for development of the *Pik3r1* mutation, and, importantly, we demonstrate that this novel SHORT syndrome model reproduces the unexplained uncoupling of metabolic dyslipidemia from lipodystrophic insulin resistance seen in humans, although the most widely studied lipogenic transcriptional programmes are only mildly perturbed.

## RESULTS

### Growth and development of Pik3r1 Y657X knockin mice

The truncating *Pik3r1* Y657X mutation, previously associated with normolipidemic severe insulin resistance in SHORT syndrome (25), was knocked into murine embryonic stem cells by homologous recombination-based gene targeting, and these cells were used to generate founder heterozygous knock-in mice (**Supplemental Figure S1**). Immunoblotting of insulin-responsive liver, skeletal muscle and adipose tissue confirmed the presence of a truncated p85α gene product, which in general was more highly expressed than full length, wild type protein in the same tissues, most likely due to loss of a previously described C-terminal ubiquitylation motif (27) No change in expression was seen of the p85β regulatory subunit of PI3K, while expression of the p110a catalytic subunit was reduced in white adipose tissue, and expression of the p110β catalytic subunit was reduced in both liver and white adipose tissue. p110β expression very low in skeletal muscle in both wild-type and mutant animals (**Supplemental Figure S1**).

Homozygosity for the *Pik3r1* Y657X allele was lethal *in utero,* with no homozygous embryos identified beyond E11.5. At E11.5 homozygous embryos were smaller, with poorly developed limb buds and reduced eye pigmentation (**Figure 1A**), while heterozygous embryos were smaller from E15.5 onwards (**Supplemental Figure S2**). No difference in mass of wild type and heterozygous placentas examined from wild type dams was seen, however vascularisation of the placental exchange region was severely compromised, with around a 40% reduction in vessel density, volume and length at E15.5 (Supplemental Figure S3), as reported previously for placentas heterozygous for a kinase dead p110α catalytic subunit (28). In contrast to that model, however, the size of the placental exchange region (both volume and surface area) and the thickness of the diffusion barrier were normal (**Supplemental Figure S3**).

**Figure 1.**
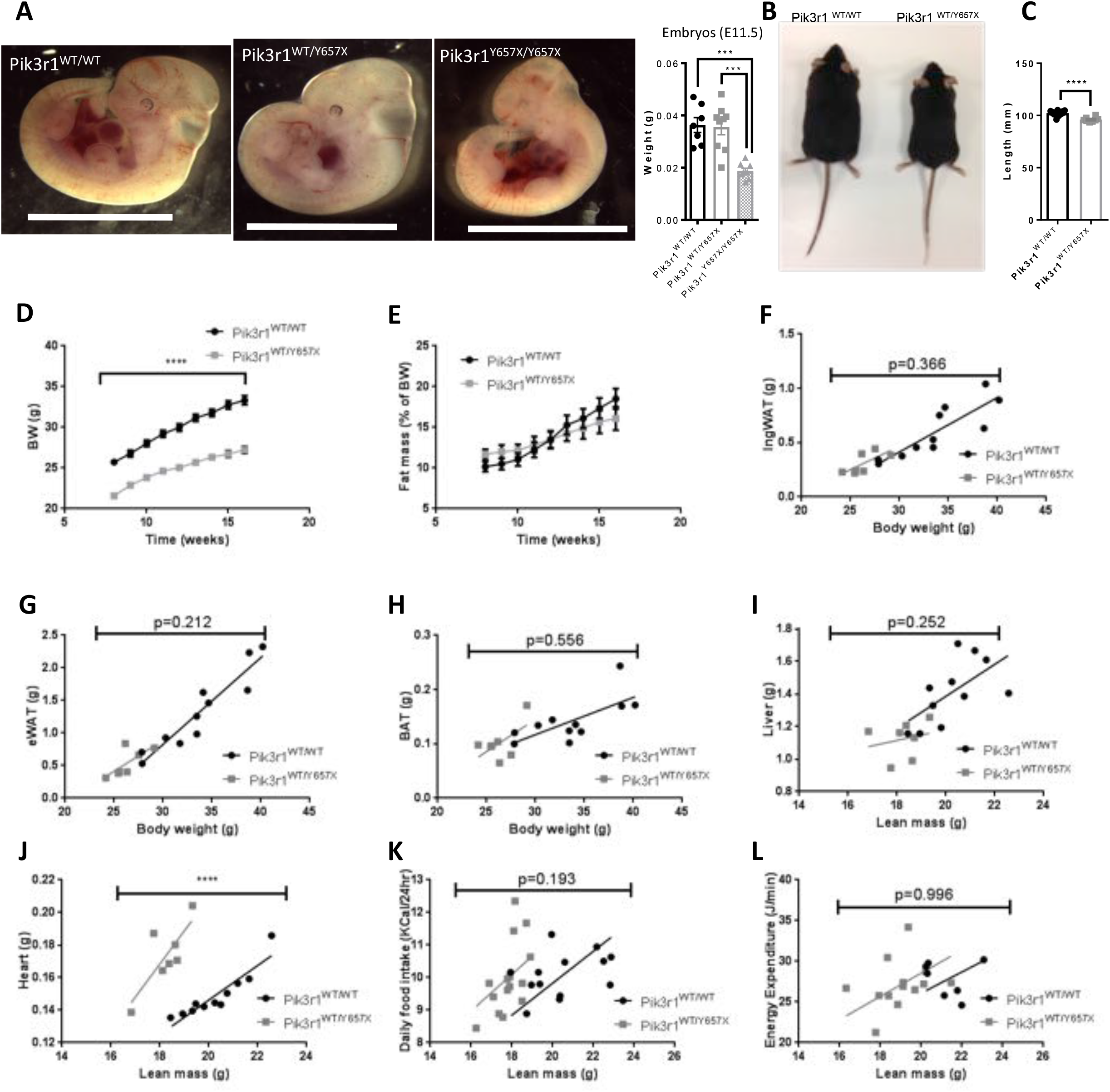
Effect of *Pik3r1* Y657X on prenatal development and postnatal growth. (A) Representative images and weights (adjacent scatter plot) of *Pik3r1^WT/WT^*, *Pik3r1^WT/Y657X^* and *Pik3r1^Y657X/Y657X^* embryos at E11.5. Scale bar = 5mm (B) Representative image of male *Pik3r1^WT/WT^* and *Pik3r1^WT/Y657X^* mice at 18 weeks old. (C) Body lengths (nose to anus) at 18 weeks of *Pik3r1^WT/WT^* and *Pik3r1^WT/Y657X^* mice (n=11 and 18 respectively) (D) Bodyweight increase from 8 to 16 weeks of *Pik3r1^WT/WT^* and *Pik3r1^WT/Y657X^* (n=16 and 12 respectively). (E) Time course of fat mass accretion, expressed as percentage of bodyweight, of *Pik3r1^WT/WT^* and *Pik3r1^WT/Y657X^* mice (n =16 and 12 respectively). Mean ± S.E.M. are shown for scatter plots in (A) and (C)-(E) (F)-(J) Masses of (F) inguinal adipose tissue (IngWAT) (G) Epididymal adipose tissue (eWAT), (H) Brown adipose tissue (BAT), (I) Liver, and (J) Heart of *Pik3r1^WT/WT^* and *Pik3r1^WT/Y657X^* mice (n= 11 and 7 respectively). (K) Food intake (n=13 for *Pik3r1^WT/WT^* and n=14 for *Pik3r1^WT/Y657X^)* and (L) Energy expenditure (n=7 for *Pik3r1^WT/WT^* and n=13 for *Pik3r1^WT/Y657X^*) of wild type and heterozygous mice assessed at 18 weeks old. All masses and energy expenditure are shown relative to total lean mass, and were analysed statistically by ANCOVA. **** = p < 0.0001

*Pik3r1^WT/Y657X^* mice were born at expected Mendelian frequency, however they showed impaired linear growth on a chow diet, with body length of males 6% less than wild-type littermates at 18 weeks old (**Figures 1B,C**), and bodyweight 17% less (**Figure 1D**). Overall body composition assessed by TD-NMR showed no difference between heterozygous males and controls (**Figure 1E**), and, in keeping with this, no difference was found in epididymal or inguinal white adipose depot nor interscapular brown adipose depot weights when the reduced bodyweight of the male heterozygous mice was taken into account (**Figure 1F-H**). Plasma leptin concentrations were similar between male *Pik3r1^WT/Y657X^* and wild-type mice (3.3±1.2 *vs* 2.6±0.3 μg/L in the fasting state (n=9,10; not significant), and 10.3±2.2 *vs* 15.9±3.9 μg/L on *ad libitum* feeding (n=13,13; not significant)). Liver weights were also indistinguishable (**Figure 1I**), however hearts were significantly heavier in *Pik3r1^WT/Y657X^* mice (**Figure 1J**) whether analysed with respect to lean mass or whole body weight by ANCOVA. No significant difference in either food consumption or energy expenditure was seen at 16 weeks old on chow diet (**Figure 1K,L**). A similar pattern of differences was seen between heterozygous and wild-type female mice for all variables assessed in both sexes (**Supplemental Figure S4**).

### Reproductive function of *Pik3r1^WT/Y657X^* mice

Postpubertal women with SHORT syndrome commonly exhibit anovulation with severe ovarian hyperproduction of testosterone (25). This is most likely a consequence of systemic severe insulin resistance, however important roles for PI3K in the ovary are also known, with deletion of PTEN, a negative regulator of PI3K, in oocytes in mice producing unrestrained maturation of primordial follicles and thus rapid exhaustion of the follicular pool (29). We thus assessed ovarian morphology and reproductive function in the *Pik3r1^WT/Y657X^* mice. No significant difference was seen in litter frequency for any pairwise genotype combination, assessed between 10 and 18 weeks, however litter size was reduced significantly for *Pik3r1^WT/Y657X^* mothers whether mating with wild-type or heterozygous males, and taking into account the reduced litter size expected in double heterozygous matings due to homozygote non-viability (**Supplemental Table S1**). There was also a surprising excess of heterozygous offspring both in double heterozygous crosses and when make heterozygotes were crossed with wil-type females, but notwhen wild-type males were crossed with heterozygous females. As expected, corpora lutea were observed in both *Pik3r1^WT/Y657X^* and wild-type females of reproductive age (at 42 days old), consistent with ovulation, and serum testosterone concentrations were similar in both genotypes at 12 weeks old (0.18 ± 0.05 μg/l in heterozygotes (n=7) *vs* 0.23 ± 0.04 in wild-types (n=10) (Not significant)). More detailed morphometric analysis of ovaries at 7 days old, before sexual maturation, showed a grossly normal microscopic appearance, with the proportions and sizes of follicles at different developmental stages similar in wild type and heterozygous ovaries, suggesting normal early follicle development (**Figure 2**).

**Figure 2.**
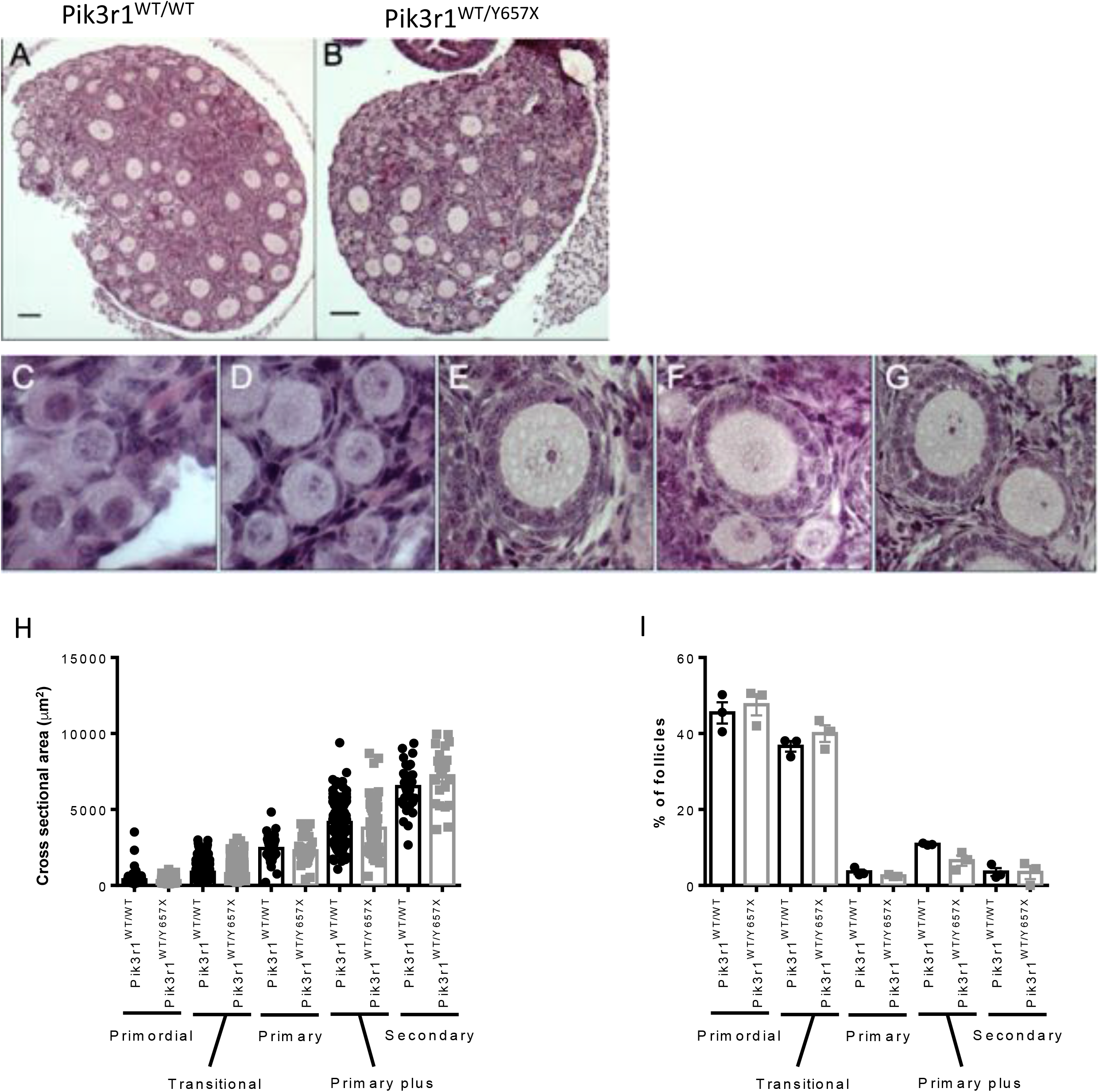
Ovarian follicular appearance and profile of *Pk3r1^WT/Y65TX^* mice and wild type littermates. Representative H&E-stained cross sections of ovaries from 7 day old *Pik3r1^WT/WT^* (A) and *Pik3r1^WT/Y657X^* (B) mice. Scale bars = 300μm. (C-G) Examples of follicles at different stages of pre-antral follicular development: primordial (C), transitional (D), primary (E), primary plus (F) and secondary (G). (H) Cross sectional area of follicles from 7 day old *Pik3r1^WT/WT^* and *Pik3r1^WT/Y657X^* mouse ovaries, classified by developmental stage (n=3 per genotype). (I) Proportions of follicles at different developmental stages in *Pik3r1^WT/WT^* and *Pik3r1^WT/Y657X^* mouse ovaries (n=3 per genotype). Bars represent mean ± SEM.

### Adipose response of *Pik3r1^WT/Y657X^* mice to a high fat diet

Lipodystrophy, or impaired accretion of adipose tissue, is a common feature of SHORT syndrome (24), and the previously reported *Pik3r1^WT/R649W^* mouse was also shown to have reduced subcutaneous adipose tissue even on chow diet. On the other hand at least one human proband with the Y657X mutation showed normal adipose development (25). To assess whether placing a greater load on adipose tissue storage in *Pik3r1^WT/Y657X^* mice would unmask a lipodystrophic phenotype, mice were fed a palatable 45% high fat diet (HFD) from 8 weeks old. *Pik3r1^WT/Y657X^* mice showed significantly reduced gain of weight and whole body fat content under these conditions over the 8 week period studied (**Figure 3A,B**). Epididymal white adipose tissue was most severely affected (**Figure 3C**), with the difference in inguinal adipose tissue not achieving statistical significance by ANCOVA (**Figure 3D**). Brown adipose tissue mass was unaffected (**Figure 3E**). Consistent with reduced adiposity, *Pik3r1^WT/Y657X^* mice had lower serum leptin concentration in the fed state (37.7±7.3 *vs* 11.3±2.0 μg/L (n=15,11; p=3x10^-3^)), although this was not significant on fasting (15.9±3.9 *vs* 10.3±2.2 μg/L (n=13,13; not significant)). Lean mass did not increase at a greater rate in *Pik3r1^WT/Y657X^* mice than wild-type littermates (**Supplemental Figure S5**), and liver weights also did not diverge, while heart weights were elevated in heterozygotes, as on chow (**Figure 3F,G**).

**Figure 3.**
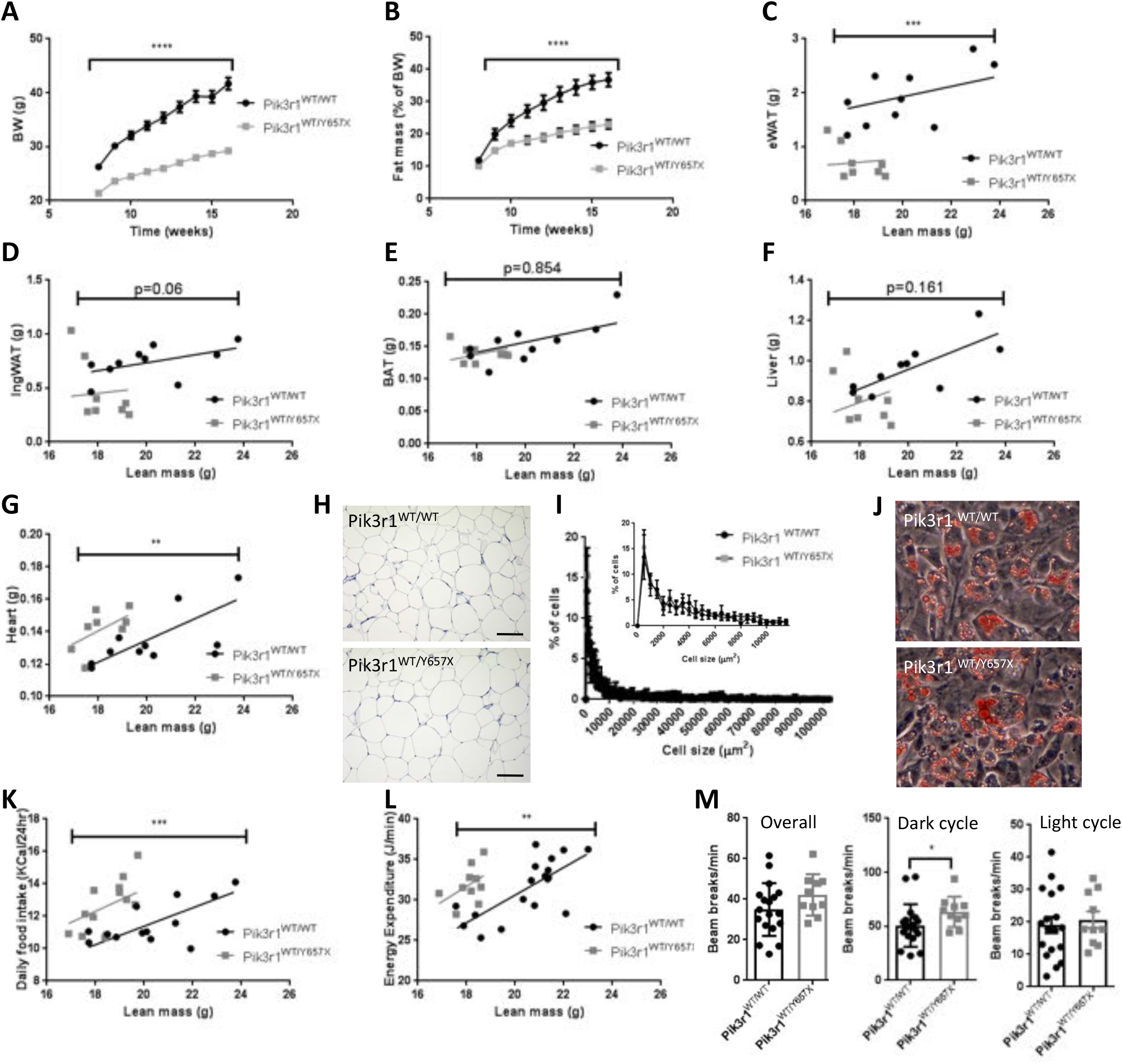
Response of *Pik3r1^WT/Y657X^* mice to a palatable 45% fat diet. (A) Bodyweight increase from 8 to 16 weeks of *Pik3r1^WT/WT^* and *Pik3r1^WT/Y657X^* (n=16 and 12 respectively). (B) Time course of fat mass accretion, expressed as percentage of bodyweight, of *Pik3r1^WT/WT^* and *Pik3r1^WT/Y657X^* mice (n =16 and 12 respectively). Masses of (C) Epididymal adipose tissue (eWAT), (D) Inguinal adipose tissue (IngWAT), (E) Brown adipose tissue (BAT), (F) Liver, and (G) Heart of *Pik3r1^WT/WT^* and *Pik3r1^WT/Y657X^* mice (n= 12 for both genotypes). (H) Representative histological appearance of haematoxylin and eosin-stained eWAT from *Pik3r1^WT/WT^* and *Pik3r1^WT/Y657X^* mice. Scale bars = 100μm (I) Adipocyte size distribution in eWAT based on quantification of >1,000 cells per genotype from 4 wild type and 4 heterozygous mice. The inset shows a zoomed-in view of the early part of the distribution. (J) Representative images of *ex vivo* differentiated stromovascular cells from ingWAT stained with Oil Red O (see also Supplementary Figure S5). (K) Food intake (n=13 for *Pik3r1^WT/WT^* and n=14 for *Pik3r1^W?/ĩ657X^)* and (L) Energy expenditure (n=17 for *Pik3r1^WT/WT^* and n=10 for *Pik3r1^WT/Y657X^)* of wild type and heterozygous mice assessed at 18 weeks old. All masses and energy expenditure are shown relative to total lean mass, and were analysed statistically by ANCOVA. **** = p < 0.0001 (M) Locomotor activity of *Pik3r1^WT/WT^* and *Pik3r1^WT/Y657X^* mice (n=17 and n-10 respectively) Mean ± SEM are shown. Statistical analysis used included 2-way repeated measured ANOVA for (A-B), ANCOVA for (C-G, K and L), and Student’s t-test for (M). *p < 0.05, **p < 0.01, ***p < 0.001 and ****p < 0.0001.

Despite the marked decrease in weight of epididymal white adipose tissue, the histological appearance and adipocyte size distribution was similar in both genotypes (**Figure 3H,I**), indicating a smaller number of normal sized adipocytes in heterozygotes. *Ex vivo* differentiation of preadipocytes from either inguinal or epididymal depots using conventional protocols was normal, however (**Figure 3J and Supplemental Figure S6**) as previously reported for *Pik3r1^WT/R649W^* mice, suggesting that the reduced adipose expansion is not accounted for by a cell autonomous defect in differentiation capacity. To assess whether the impaired adipose expansion might instead reflect altered energy homeostasis, food intake and energy expenditure were next assessed. *Pik3r1^WT/Y657X^* mice, at odds with the observed reduction in white adipose tissue expansion on high fat feeding, exhibited significantly increased food intake (**Figure 3K**). Energy expenditure was also increased, however (**Figure 3L**), possibly in part due to increased locomotor activity during the dark cycle, although observed activity was not significantly increased across the whole 24 hour cycle (**Figure 3M**).

### *Pik3r1^WT/Y657X^* mice are severely insulin resistant

In accord with the critical role known to be played by PI3K in insulin action, severe insulin resistance is common in SHORT syndrome, and insulin resistance was also reported in *Pik3r1^WT/R649W^* mice. Assessed at 12 weeks old, neither male nor female *Pik3r1^WT/Y657X^* mice on chow were hyperglycemic compared to wild-type littermates, but plasma insulin concentrations were significantly raised, more severely so in the fed state, as in human severe insulin resistance (**Table 1, Supplemental Table S2**). On high fat feeding male *Pik3r1^WT/Y657X^* mice remained hyperinsulinemic, but the difference between mutant and control animals was abolished as wild-type mice exhibited a much greater increase in insulin concentrations on high fat feeding than *Pik3r1^WT/Y657X^* animals. Glucose concentrations were slightly lower in the fed state on high fat diet in the heterozygous mice than controls, which may also have contributed to the loss of a significant difference in insulin concentrations. Plasma adiponectin concentrations in fasted chow-fed *Pik3r1^WT/Y657X^* mice were lower than in wild-type controls, however no significant difference in concentrations were seen between genotypes in the other conditions despite severe insulin resistance of heterozygous mice (**Table 1**).

**Table 1.**
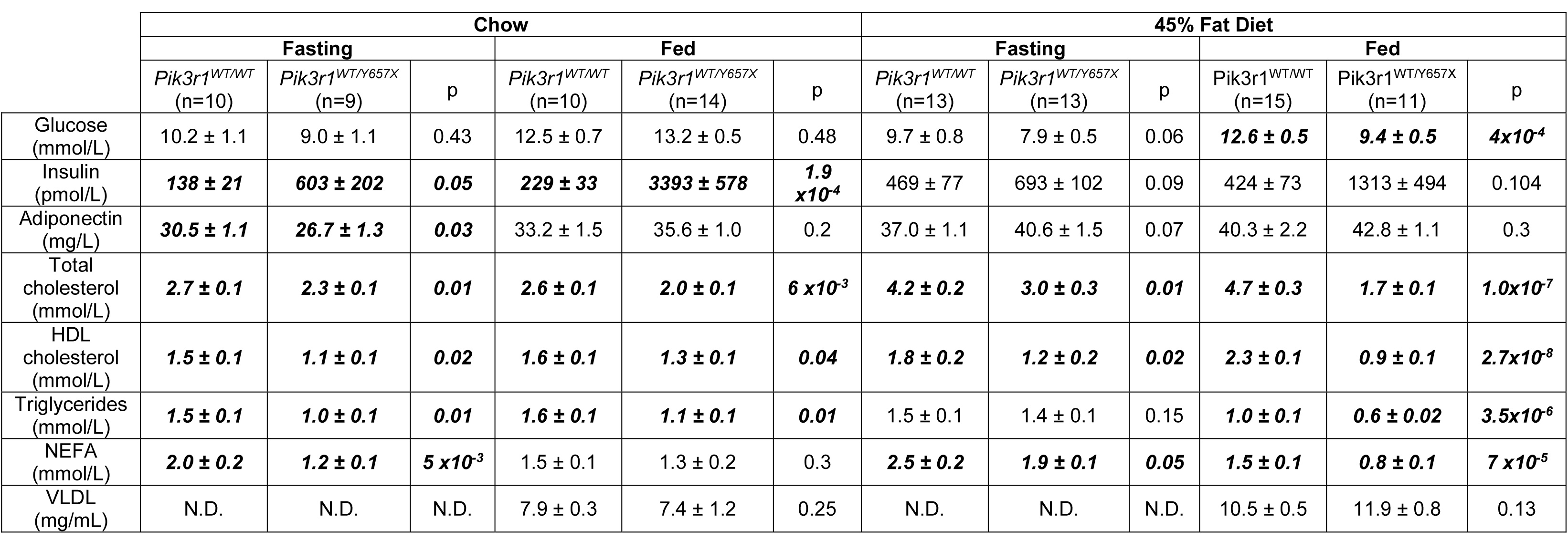
Fasting plasma biochemical profile of male *Pik3r1^WT/WT^* and *Pik3r1^WT/Y657×^* mice. Significant differences between genotypes are shown in bold. See also Supplemental Table S1 for fasting biochemical profile for chow-fed female mice. N.D. = not determined. Statistical comparisons were undertaken using the student’s t-test.

Moreover in *Pik3r1^WT/Y657X^* animals on HFD, adiponectin normalised to body fat content was significantly higher than in controls (**Supplemental Figure S7**). Normalised plasma leptin concentrations, in contrast, were lower in heterozygous mice than controls (**Supplemental Figure S7**).

To evaluate insulin sensitivity in more detail, hyperinsulinemic euglycemic clamps were undertaken on both chow-fed or HFD-fed male mice. Initial clamps undertaken without isotopic tracers demonstrated that on chow feeding *Pik3r1^WT/Y657X^* mice required a glucose infusion rate at steady state of only one eighth of the rate required by wild-type mice, confirming severe insulin resistance (**Figure 4A**). On HFD *Pik3r1^WT/Y657X^* mice remained extremely insulin resistant, however the difference between heterozygous and wild-type mice was much smaller than on chow due to insulin resistance induced in wild-type animals (**Figure 4A**). Further studies were thus undertaken on chow. Glucose and insulin excursions on oral glucose tolerance testing showed only a trend towards an increase in male *Pik3r1^WT/Y657X^* mice (**Figure 4B,C**), but were significantly increased in female mice (**Supplemental Figure S8**), while the hypoglycemic response to insulin was clearly attenuated on insulin tolerance testing in both sexes (**Figure 4D and Supplemental Figure S8**). Further clamp studies in males incorporating isotopic tracers showed glucose disposal to be 19% lower in *Pik3r1^WT/Y657X^* mice (**Figure 4E**) while suppression of hepatic glucose production by hyperinsulinemia was 49% lower compared to wild type littermates (**Figure 4F**). Despite this, liver glycogen content was similar between heterozygous *Pik3r1^WT/Y657X^* mice and wild type littermates in both fed and fasted states (**Figure 4G**). Insulin infusion lowered plasma free fatty acid concentration in plasma by 38% in wild-type mice, however this suppression was nearly abolished in heterozygous mutant animals, consistent with impaired suppression of triglyceride lipolysis in adipose tissue (**Figure 4H**).

**Figure 4.**
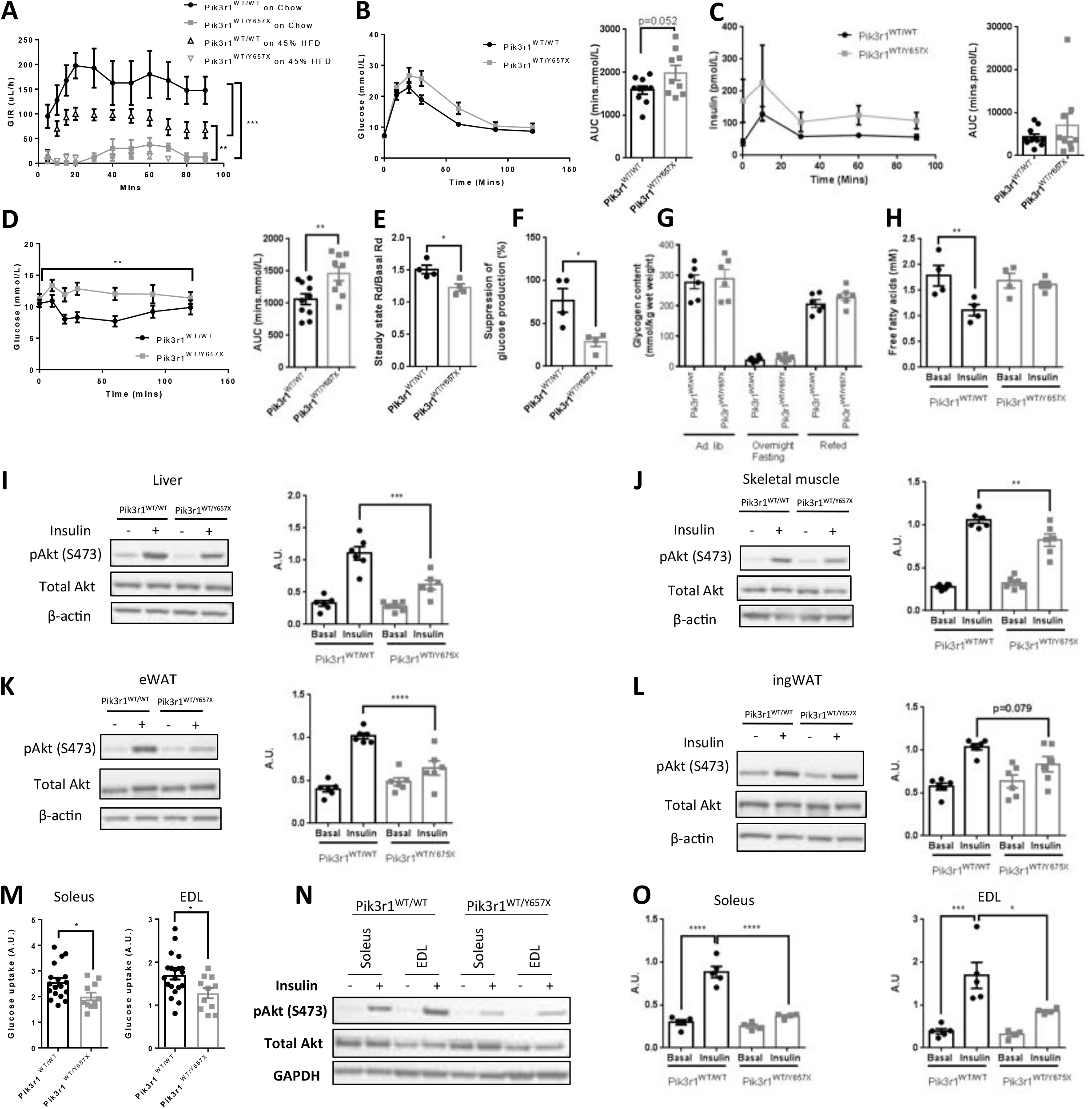
*Pik3r1^WT/Y657X^* mice show systemic and tissue-level insulin resistance. (A) Glucose infusion rates during hyperinsulinemic euglycemic clamping of *Pik3r1^WT/WT^* and *Pik3r1^WT/Y657X^* mice on chow (n=4 and 4) or 45% fat diet (n=10 and 11) at 16 weeks old; (B) Oral glucose tolerance test (OGTT) and corresponding comparison of areas under the curves (AUC) of *Pik3r1^WT/WT^* and *Pik3r1^WT/Y657X^* mice on chow at X weeks old (n=10 and 9). (C) Insulin concentrations and AUC for the same OGTT. (D) Insulin tolerance test and AUC comparison for the same mice 1 week later. (E) Glucose disposal and (F) suppression of hepatic glucose output by insulin during hyperinsulinemic euglycemic clamping of *Pik3r1^WT/WT^* and *Pik3r1^WT/Y657X^* mice on chow at 18 weeks old (both n=4). (G) Glycogen content of livers during a fasting-refeeding cycle in chow fed animals at 16 weeks old (both n=6). (H) Plasma non-esterified free fatty acid concentrations during hyperinsulinemic euglycemic clamping (both genotypes n=4). (I)-(L) Representative images of immunoblots and corresponding quantifications of tissue lysates from mice injected intraperitoneally with 2U/kg insulin 10 mins prior to sacrifice, showing pAkt^Ser473^, total Akt and their ratio: (I) Liver, (J) Skeletal muscle (K) eWAT, (L) ingWAT. (n=6 per genotype and condition). (M) Insulin-induced fold increase of glucose uptake into *ex vivo* incubated soleus (n=18 for *Pik3r1^WT/WT^,* n=11 for *Pik3r1^WT/Y657X^)* and Extensor Digitorum Longus (EDL) (n=20 and 11). (N) Representative immunoblots of Soleus and EDL lysates from the same paradigm. (O) Quantification of pAkt ^Ser473^ to total Akt ratios from soleus and EDL immunblots (n=5 and 4 for both). Quantitative data are presented as mean ± SEM. Analysis used includes 2-way repeated measures ANOVA (A-D, when 2-factors involved), Student’s t-test for (B-H, M-O) and 1-way ANOVA for (I-L). * = p < 0.05, ** = p < 0.01, *** = p < 0.001 and **** = p < 0.0001.

Intraperitoneal injection of insulin in male mice strongly induced Akt Ser 473 phosphorylation in liver, skeletal muscle and both epididymal and inguinal adipose tissue of wild-type controls, as expected, and this was reduced by 55%, 20%, 37% and 30% respectively in male *Pik3r1^WT/Y657X^* mice (**Figure 4I-L**). A similar pattern was seen for threonine 308 phosphorylation (**Supplemental Figure S9**). As muscle insulin sensitivity was previously reported not to be reduced in *Pik3r1^WT/R649W^* mice, insulin responsiveness of soleus and extensor digitorum longus (EDL) muscles was further assessed *ex vivo.* Soleus and EDL muscle from *Pik3r1^WT/Y657X^* mice both showed a 1.3-fold reduction in insulin-stimulated deoxyglucose uptake compared to wild-type muscle (**Figure 4M**). Insulin-induced Akt phosphorylation was also markedly reduced in both types of muscle *ex vivo* to a much more significant degree than *in vivo* (**Figure 4N,O**).

### *Pik3r1^WT/Y657X^* mice are hypolipidemic

Having shown that, like humans with SHORT syndrome, heterozygous *Pik3r1^WT/Y657X^* mice show severe insulin resistance and lipodystrophy, we next assessed whether, like humans, they are protected from fatty liver and metabolic dyslipidaemia. Strikingly, in both fed and fasting states, and on either chow or high fat diets *Pik3r1^WT/Y657X^* mice showed hypolipidemia, with lower plasma total cholesterol and HDL cholesterol (**Table 1**). Plasma triglyceride concentration was also lower in heterozygous animals in all cases except the fasting state after high fat feeding, and plasma free fatty acid concentration in all cases except chow fed animals in the fed state. A similar pattern of hypolipidemia was seen in chow fed female mice in the fasting state (**Table S2**). Importantly no difference was seen in immunoassay-determined VLDL concentrations in either chow fed or high fat diet fed mice (**Table 1**).

No difference was seen in liver triglyceride content between heterozygous and wild-type mice either on chow diet or on HFD, assessed both by Oil Red O staining and biochemical quantification (**Figure 5A,B**), in keeping with the lack of difference in liver weights previously noted. Liver cholesterol content was also the same in chow-fed animals of both genotypes (**Figure 5C**). To screen for abnormal lipid absorption or handling by the intestine oral lipid tolerance testing was undertaken, but although lower plasma triglyceride concentrations were seen at all time points, the excursion after lipid loading was the same in both genotypes (**Figure 5D**), and bomb calorimetry of faeces on chow showed the same energy content in both groups (**Figure 5E**). Collectively these findings argue strongly against abnormal intestinal lipid absorption and/or mobilisation as the explanation for hypolipidemia and failure to gain adipose mass on HFD.

**Figure 5.**
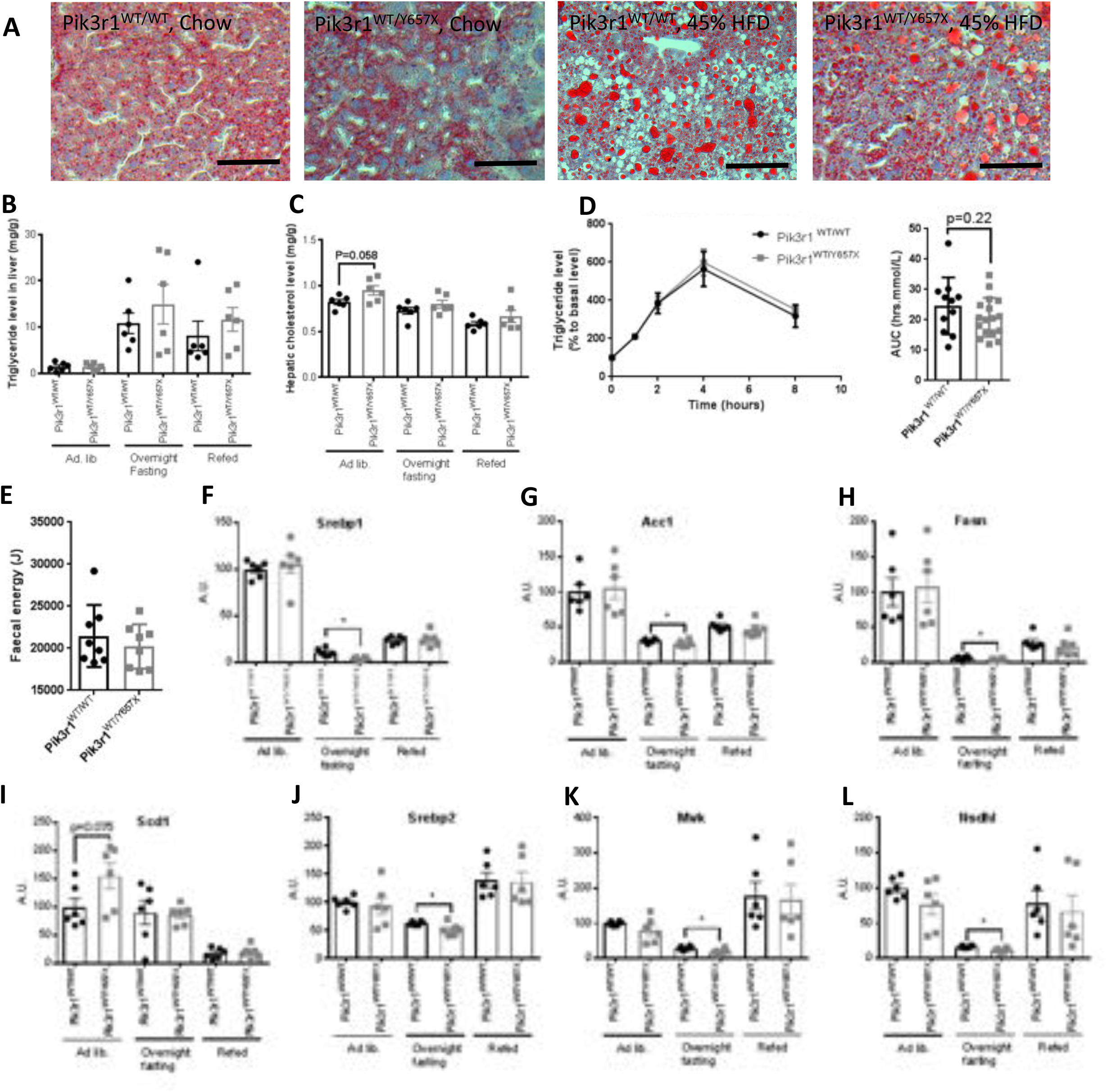
Lipid handling and liver phenotype of *Pik3r1* mice. (A) Representative images of Oil-Red-O-stained livers of chow-fed and 45% fat diet-fed *Pik3r1^WT/WT^* and *Pik3r1^WT/Y657X^* mice. Scale bars = 200μm (B) Hepatic triglyceride and (C) Hepatic total cholesterol concentration during a fasting refeeding cycle of chow fed mice at 16 weeks old (n=6 per genotype). (D) Lipid tolerance testing and comparison of areas under the curve (AUC), also of chow-fed mice, at 16 weeks old. Triglyceride concentrations were equalised at baseline by adding the difference between genotypes to the lower (heterozygous value), and the same fixed correction was applied to all points on the graph (n = 11 and n=17 for wild-type and heterozygous mice respectively) (E) Faecal energy content determined by bomb calorimetry of chow fed mice at 16 weeks old (n=8 and 8). (F)-(L) Liver mRNA expression, determined by quantitative real time PCR of (F) *Srebp1* and its transcriptional targets (G) *Acc1,* (H) *Fasn,* and (I) *Scd1,* and of (J) *Srebp2* and its transcriptional targets (K) *Mvk* and (L) *Nsdhl* in chow fed mice during a fasting refeeding cycle at 16 weeks old (n=6 per genotype per condition). Numerical data are presented as mean ± SEM, and were analysed by Student’s t-test and 2-way repeated measures ANOVA when appropriate. * = p < 0.05.

*Srebp1* and *Srebp2* are critical transcriptional regulators of *de novo* lipogenesis and cholesterol synthesis respectively, and many studies have demonstrated that they are activated by insulin in a PI3K-dependent manner. Their mRNA expression, together with expression of a panel of Srebp1 target genes *(Acc1, Fasn, Scd1)* and Srebp2 target genes *(Mvk, Nsdhl),* were assessed in the liver of chow fed animals in the fed, fasting and refed states (**Figure 5F-L**). After 16 hours of fasting there was a significant decrease in expression of all genes except *Scd1* in *Pik3r1^WT/Y657X^* mice, however no difference was seen either in the *ad libitum* fed state or after 6 hours of refeeding. Moreover expression of all genes except *Scd1* was much higher in both fed and refed states than in fasting, raising questions about the relevance of the prolonged fasting state to the observed hypolipidemic phenotype of *ad lib* fed animals.

Other candidate transcriptional regulators relevant to hypolipidemia include carbohydrate response element binding protein *(Chrebp)* and *Xbp1,* with the latter having been shown to bind p85α, facilitating its nuclear translocation (19, 20, 26) A sentinel Chrebp-responsive gene *(Pklr)* showed mild but significant repression in the fasting state, but two *Xbp1* target genes *(Acacb* and *Dgat2)* showed no transcriptional repression during a fasting refeeding cycle (**Supplementary Figure S10**). It has recently been proposed that acute insulin and Foxo-mediated modulation of the ratio between Glucose-6-phosphatase *(G6pc)* and Glucokinase (*Gck*) serves to toggle the direction of hepatocyte glucose flux between lipogenesis and hepatic glucose production, and that this mechanism is operative before transcriptional changes in canonical lipogenic transcription factors are observed (30). However although an increase in *G6pc:Gck* liver transcript levels was seen on fasting of wild-type mice in this study, consistent with this model, this ratio was suppressed in *Pik3r1^WT/Y657X^* mice, as reported in mice fed a Western-type diet (30).

## Discussion

*PIK3R1* was first sequenced in the 1990s and since then it has been intensively studied in different cellular contexts, with many different constitutional and conditional murine genetic models generated. These studies have collectively established that *PIK3R1* gene products are critically involved in signaling downstream from tyrosine kinase receptors and also that they serve to integrate this role with other signaling inputs, most commonly through interaction of relevant proteins with the N terminal domain of p85a, which is truncated in the splice variants p55α and p50α. *PIK3R1* gene products may also play signaling roles independent of RTKs.

Human pathogenic mutations in *PIK3R1* have only been discovered relatively recently. Selective loss of p85α produces agammaglobulinemic immunodeficiency (10), closely replicating findings in the earlier, corresponding murine model (9), while somatic activating mutations are found in some solid malignancies (31), in line with the role of PI3K in transducing growth factor signaling. Germline activating mutations, however, produce immunodeficiency due to impaired lymphocyte maturation (32, 33), with no consistent metabolic or growth abnormalities described. SHORT syndrome, the last disorder to be associated with *PIK3R1* mutations, is predominantly associated with mutations in the C terminal SH2 domain of p85α/p55α/p50α (21–23), and provides arguably the most surprising and yet instructive insights into *PIK3R1* function in humans, particularly in view of discordant phenotypic features between it and the various murine *Pik3r1* knockout models that have been reported.

Another mouse model of SHORT syndrome has recently been described, heterozygous for the common SHORT-associated R649W mutation in the phosphotyrosine binding motif of the C terminal SH2 domain (26). This model recapitulated key features of SHORT syndrome including growth impairment, lipodystrophy and insulin resistance. While the phenotype of the novel model we describe is broadly concordant this model, it differs in some significant respects, and we moreover extend phenotyping with more detailed developmental assessment and challenge with a high fat diet.

As previously described for the R649W mutation, no mice homozygous for the *Pik3r1* Y657X mutation were born, contrasting with *Pik3r1* knockout mice which are born but die perinatally (9). *Pik3r1^Y657X/Y657X^* embryos are not seen beyond E11.5, at which stage they are smaller and retarded in development compared to wild-type and heterozygous embryos. This timing is reminiscent of mice with knockout of both *Pik3r1* and *Pik3r2,* which do not survive beyond E12.5 (15). Homozygous knockout or homozygosity for a kinase dead mutant of p110α also both lead to lethality around E10.5 (5, 34), while embryos without functional p110δ are viable (35), and those without functional p110β have been reported to have variable phenotypes (8). These findings demonstrate that the SHORT-related allele is not simply a loss-of-function allele, but that instead it disrupts *Pik3r1* function in a manner that renders it resistant to rescue by other PI3K regulatory subunits. Pertinent to this are prior findings that expression of p85β is upregulated and expression of all three p110 catalytic subunits downregulated in cells from p85α knockout mice (9, 36, 37). In *Pik3r1^WT/Y657X^* mice, as in dermal fibroblasts from humans with SHORT syndrome (25), p85β expression is unaffected, however p110α was modestly downregulated in subcutaneous adipose tissue only, and p110β more markedly downregulated in both white adipose tissue and liver. These findings suggest that it may be the failure to upregulate p85β that is most important in the phenotypic differences between p85a knockout and *Pik3r1^WT/Y657X^* mice.

Reduced embryo size may be due to an embryo autonomous growth defect, to impaired placental function, and/or to perturbed placental nutrient flux. Indeed, recent studies of mice heterozygous for a kinase dead p110α allele *(Pik3ca^KD/WT^)* have shown that p110a action on both maternal and fetal sides of the placental circulation influence each of these (28). *Pik3r1^WT/Y657X^* fetuses are around 20% smaller that wild-type fetuses at E15.5, which is similar to the decrease noted in *Pik3ca^KD/WT^* fetuses previously. Although no significant reduction in placental mass was seen, striking reduction of the vascularisation of the placental exchange region was seen at E15.5, again very similar to the *Pik3ca^KD/WT^* model. In contrast to that model, however, the size of the placental exchange region and thickness of the diffusion barrier were normal for *Pik3r1^WT/Y657X^* fetuses. These findings are consistent with p85α and p110α co-operating in formation of the placental vascular tree, and add to prior evidence highlighting the importance of class IA PI3Ks (particularly p110a) in developmental angiogenesis (38).

In the previous study of *Pik3ca^KD/WT^* mice, maternal genotype as well as fetoplacental genotype was important in determining placental phenotype. This was not assessed directly in this study, where wild-type dams were used for developmental studies, however a marked effect of maternal genotype was seen on litter size, which were reduced by around 50% of the expected for heterozygous dams, and apart from homozygous lethality no evidence of selection for fetal genotype was apparent. Whether this relates to reduced size of the heterozygous dams, to their metabolic state during gestation, or both, remains to be determined. As previous studies have suggested that the insulin resistance of pregnancy is caused by increased *Pik3r1* expression (39), it will be interesting to assess whether pregnancy exaggerates the already severe insulin resistance seen in *Pik3r1^WT/Y657X^* mice in the non pregnant state, and whether any such exacerbation influences growth of the fetoplacental unit.

PI3K also plays several important roles in ovarian function, some of which, such as the activation of dormant primordial follicles, are follicle autonomous and some of which are indirect, mediated by factors such as systemic insulin resistance. This is manifest in the severe hyperandrogenism and ovulatory dysfunction seen in young women with SHORT syndrome (25). Despite these well established links, we report no evidence of altered early follicle development, nor evidence of impaired fertility nor hyperandrogenism in female *Pik3r1^WT/Y657X^* mice. This suggests either that *Pik3r1* gene products are not involved in triggering oocyte maturation, or that the degree of PI3K hypofunction is insufficient to compromise this action. The lack of features of insulin resistance-related ovulatory dysfunction and hyperandrogenism is in keeping with many prior studies showing that mice model this highly prevalent aspect of human severe insulin resistance poorly (40).

SHORT syndrome commonly features partial lipodystrophy (24, 25), and on chow diet the previously reported *Pik3r1^WT/R649W^* mouse, too, was reported to have reduced subcutaneous adipose depots (26). In this study no significant differences in whole body fat content and adipose depot weights were observed on chow when the smaller size of the *Pik3r1^WT/R649W^* mice was taken into account, however they diverged strikingly between wild type and heterozygotes on high fat feeding. *Ex vivo* differentiation of white preadipocytes was not impaired, consistent with the previous study, although in preadipocyte cell lines overexpression of the mutant has been shown to impair adipogenesis markedly (25). It was previously suggested that a reduced number of preadipocytes in adipose tissue might account for reduced adipose expansion, however although this was not assessed in the current study, the normal adipocyte size and histological appearance argue against this, as a larger adipocyte size would be expected if a smaller number of precursor cells were accommodating the same hypercaloric load. Alternative explanations for reduced adipose accretion that are disproven by the current study are reduced food intake or intestinal lipid malabsorption, or increased partitioning of excess energy into growth of non adipose tissues.

Lipodystrophy in humans and mice is normally associated strongly with exaggerated features of obesity-associated metabolic syndrome including fatty liver, metabolic dyslipidaemia, and low plasma adiponectin concentrations (41, 42), however SHORT syndrome is unique among human insulin resistant partial lipodystrophies in featuring reduced adiposity with preserved plasma adiponectin levels, absence of fatty liver and apparent protection from metabolic dyslipidemia (24, 25). This is similar to the profile seen in people with severe insulin resistance due to mutations in the insulin receptor gene (43), suggesting that proximal insulin signalling defects in humans produce a distinct subphenotype of insulin resistance. The mechanisms underlying these human observations have not been tested, however insulin resistance has been uncoupled from hyperlipidemia in various mouse models, including mice with liver-specific knockout of *Insr* (44), both *Irs1* and *Irs2* (45), *Pik3ca* (46) or *Akt2* (47). These severe perturbations have generally been suggested to produce hypolipidemia through loss of insulin-stimulated upregulation of genes involved in *de novo* lipogenesis and/or cholesterol synthesis. The recent *Pik3r1^WT/R649W^* SHORT syndrome model mice, in contrast, were said to have normal blood lipid concentrations, however in the case of the *Pik3r1^WT/Y657X^* mouse we report hypolipidemia in both males and females, on chow diet and high fat feeding, and in both the fed and fasted state. Moreover liver triglyceride levels showed no difference between wild-type and *Pik3r1^WT/Y657X^* mice, although in the face of lipodystrophic severe insulin resistance they would normally be increased. This suggests that this murine model may permit dissection of mechanisms underlying the dyslipidemia-insulin resistance dissociation.

Preliminary assessment of lipogenic transcriptional effects of the *Pik3r1* Y657X mutation in a feed/fasting/refeeding paradigm did show significant reduction in transcript levels of *Srebp* and their transcriptional targets in the liver, with a sentinel CΛrebp-responsive gene also mildly suppressed, however these changes were only seen in the fasting state. Whether these mild fasting changes can explain the hypolipidemia observed, which is most striking in the fed state on high fat diet, is unclear. No evidence was found for reduced lipogenic transcriptional activity of *Xbp1,* which has both been shown to interact with p85α (18, 19), and to serve as a potent lipogenic transcription factor (20). A further mechanism toggling glucose routing between hepatic lipogenesis and glucose secretion depends on the relative expression of glucose-6-phosphatase *(G6pc)* and glucokinase (Gck), with a high G6pc:Gck ratio favouring hepatic glucose output and disfavouring lipogenesis (30). However although the refed *G6pc:Gck* ratio did significantly differ in *Pik3r1^WT/Y657X^* mice compared to controls, the direction of the difference would be predicted to favour lipogenesis.

Interestingly, global knockout of the insulin-responsive *Glut4* transporter in mice grossly mimics the phenotype of the mice we describe in key respects, featuring insulin resistance, severely reduced adipose tissue, and an enlarged heart (48). Moreover, although fatty liver was not observed, blood lipid levels and liver triglyceride secretion was markedly increased (49), but was countered by accelerated VLDL clearance by peripheral tissues, demonstrating extensive remodelling of metabolism (50). In this case lipid tolerance testing does not provide evidence for similarly enhanced clearance of lipid, however. Gaining further insight into the mechanisms uncoupling dyslipidemia from insulin resistance in *Pik3r1^WT/Y657X^* mice will require metabolic flux studies, and consideration of the possibility that effects on hepatic metabolism are effected by primary dysfunction in other insulin sensitive organs such as adipose tissue, which has recently been shown in some contexts to be critical for such indirect regulation ofhepatic gluconeogenesis (51, 52).

In summary, we describe a novel mouse model of SHORT syndrome driven by a naturally occurring human mutation in *PIK3R1.* We describe novel developmental, reproductive and metabolic aspects of the model, and establish that it recapitulates the highly unusual human observation of lipodystrophic insulin resistance uncoupled from atherogenic dyslipidaemia.

## METHODS

### Mice generation and maintenance

A vector containing the *Pik3r1* exon 15 Y657X mutation and a neomycin resistance cassette flanked by LoxP sites was introduced into the genome of C57Bl/6 embryonic stem cells by electroporation and homologous recombination. Targeted cells were injected into Bl/6J blastocysts to generate chimeras which were bred onto the C57Bl/6J background to create the mutant strain (**Supplemental Figure S1**). All animals were kept on a C57Bl/6J background, backcrossed at least three times, and housed on a 12-hour-light/12-hour-dark cycle at 23°C, with *ad libitum* access to water and food. Feed was with either chow (Catalogue no. 105, Safe diets) or a 45% high fat diet (45% fat, 35% Carbohydrate and 25% protein) (Catalogue no. D12451, Research diet, Inc). All animal experiments were carried out under the UK Home Office Animals (Scientific Procedures) Act 1986, following ethical review by the University of Cambridge.

### Fetoplacental development studies

Single male mice were housed with a female mouse and noon of the day of detection of a vaginal plug was taken to be E0.5. Images of dissected embryos were acquired with a Zeiss SteREO Discovery V8 microscope and AxioCam MRc5 camera. Placentas were bisected and one half was fixed in 4% (wt/vol) paraformaldehyde, paraffin-embedded, exhaustively sectioned at 7 μm and stained with hematoxylin and eosin to analyse gross placental structure. The other half was fixed in 4% (wt/vol) glutaraldehyde, embedded in Spurr’s epoxy resin and a single, and 1 μm midline section was cut and then stained with toluidine blue for detailed analysis of labyrinthine (exchange) zone structure using the Computer Assisted Stereological Toolbox (CAST v2.0) program as previously described (28).

### Assessment of growth, body composition, and energy homeostasis

Fat and lean mass of live mice was determined by time-domain NMR using a Bruker’s minispec whole body composition analyser. For determination of food intake and energy expenditure, male mice were acclimatized for 1 week to single housing then given 200g of standard chow diet or 45% high fat diet for 10 days. Food intake was measured every 24 hours, and energy expenditure was measured by indirect calorimetry as described previously (53). Locomoter activity was quantified as beam breaks over 48 hours. Fecal energy content was measured by drying then totally combusting feces from 48hr calorimeter runs in an IKA Calorimeters Oxygen Bomb calorimeter (IKA C1).

### Biochemical assays

Blood was collected by cardiac puncture and plasma separated and snap frozen immediately. Plasma analytes were assayed following manufacturers’ instructions with the following assays: glucose (Siemens Healthcare Diagnostics), insulin (Meso Scale Discovery), leptin (Meso Scale Discovery), adiponectin (Meso Scale Discovery), triacylglyceride (Siemens Healthcare Diagnostics), total cholesterol (Siemens Healthcare Diagnostics), HDL cholesterol (Randox), VLDL (Cusabio), free fatty acids (Roche) and testosterone (IBL international). Plasma insulin concentration during glucose tolerance testing used an ELISA (Crystal Chem).

Hepatic glycogen content was determined in snap-frozen liver samples that were weighed before glycogen hydrolysis in 1 M HCl (2.5 h at 100°C). Hydrolysate was neutralized with NaOH and glucose units analysed fluorometrically as described in (54). Hepatic triglyceride was extracted and measured with a Triglyceride Colorimetric Assay kit (Cayman) according to the manufacturer’s instructions. Hepatic cholesterol was extracted from 10mg liver using 400uL of chloroform:isopropanol:NP-40 mixture (7:11:0.1) in a homogenizer before centrifugation and air drying of superatant at 50°C. After 30 minutes of vacuum drying lipids was dissolved in PBS and cholesterol was assayed with Siemens Dimension Healthcare following the manufacturer’s instructions (Siemens Healthcare Diagnostics).

### *In vivo* metabolic studies

For fasting/refeeding studies, mice were fed *ad libitum* on the standard chow until 16 weeks before fasting for 16 hours and refeeding with chow for 6 hours. Tissues were harvested and snap-frozen before overnight fasting, after fasting and then after 6 hour refeeding. For study of acute insulin action 16 hours fasted mice (16 weeks old) were injected with 2U/kg insulin intraperitoneally and tissues were collected after 10 minutes and snap frozen in liquid nitrogen. Glucose tolerance testing was undertaken on overnight (16 hour) fasted mice using 2g/kg 20% glucose *via* oral gavage.For insulin tolerance testing mice were fasted for 6 hours and challenged with 0.5U/kg intraperitoneal insulin. For lipid tolerance testing overnight 16 hour fasted mice were administered 10mL/kg olive oil by oral gavage.

Hyperinsulinemic euglycemic clamp studies and the subsequent sample processing and analysis were conducted as described previously (55) with minor modifications. 16 week old mice were fasted overnight, before clamps were carried out with a priming dose of human insulin (0.7 mU), followed by a constant insulin and [3-^3^H]-D-glucose tracer infusion at 7 mU/h and 0.72 μCi/h, respectively.

### Study of insulin action *ex vivo*

Measurement of insulin action on isolated Soleus and Extensor Digitorum Longus was undertaken in 16 week old male mice as previously described(56). For primary preadipocyte studies fat pads from 8 week-old male mice were isolated, minced and digested in pre-warmed sterile digest solution (1:3 BSA 7.5 % solution:Hanks’ Buffered Salt Solution, and 1 mg/ mL Collagenase Type II). Digested tissues were strained through 100 μm cell strainers and placed on ice for 20 minutes. Following removal of the fat layer, supernatant was mixed 1:1 with growth medium (high-glucose DMEM, 10% NCS, 4 mM L-glutamine and 100 U/L penicillin streptomycin), and the stromovascular fractions washed twice with growth medium, and resuspended in differentiation medium (growth media supplemented with 2.4 nM insulin and 150 μM sodium L-ascorbate). The cells were plated into two wells of a 12-well plate and fed with fresh differentiation medium daily for 3 days and then every other day until Oil-Red O staining as described previously(25).

### Histology

Ovaries from day 7, 21 or 42 (D7, D21 or D42) wildtype and heterozygous female mice (3 per age per genotype) were fixed in 10% neutral buffered formalin solution, dehydrated in ethanol, infiltrated with Histo-Clear, and embedded in paraffin. 5μm serial sections were stained with haematoxylin and eosin. Follicles in every 5^th^ section of D7 ovaries were classified morphologically as primordial, transitional, primary, primary plus or secondary, and their cross-sectional area measured in ImageJ(57). To assess ovulation, every 10^th^ section of the D42 ovaries was analysed for the presence of corpora lutea.

Adipocyte size measurement was undertaken in formalin-fixed, paraffin-embedded adipose tissue stained with haematoxylin and eosin. Cell sizes were measured using CellP software (Olympus Scientific Solutions). At least 1000 cells per genotype were analyzed. For oil-red-O staining of liver, livers were snap-frozen and embedded in optimum cutting temperature (OCT) media. Following sectioning with a cryostat, sections were fixed in 10% NBF, incubated in 0.5% oil red O working solution, and counterstained with haematoxylin. Images were taken with an Olympus CKX41 inverted microscope and Olympus DP20 microscope camera.

### Gene expression analysis

For protein expression studies, freshly dissected issues were snap-frozen in liquid nitrogen. Liver and skeletal muscle were homogenised using MD ceramic beads and an MD machine in RIPA buffer with proteinase and phosphatase inhibitors (Roche). Adipose tissue was ground using mortar and pestle in liquid nitrogen before dissolving in the buffer. Protein concentrations were determined using the BCA assay (BioRad). Western blotting employed the Novex gel system (Thermo Fisher Scientific). Antibodies used are shown in **Supplemental Table S3** Imaging and quantification of immunoblots were undertaken using the BioRad image system.

For RNA expression studies total RNA was extracted by RNeasy kits (Qiagen). 500 ng of total RNA was used for cDNA synthesis with the ImProm-II reverse transcription system (Promega). Realtime quantitative PCR was carried out in 13 μl reaction volumes containing 300 nM forward and reverse primers, 150 nM fluorogenic probe when applicable, and ABI Taqman or SYBR^®^ Green Master Mix (Thermo Fisher Scientific). Reactions were conducted in duplicate with a standard curve for each gene using the QuantStudio™ 7 Flex Real-Time PCR system (Thermo Fisher Scientific). Expression of target genes were normalised to the geometric mean of 4 housekeeping genes (*Ywhaz, Ppia, B2m* and *Eef1a1*) (see **Supplemental Table S4** for primer and probe sequences).

### Statistical Analysis

Numerical data are presented as mean ± SEM and statistical tests used are indicated in figure legends. In general unpaired two-tailed Student’s t tests were used to compare two groups of data, while ANOVA with *post hoc* testing as indicated was performed for more than two groups. All these analyses were performed using GraphPad Prism (GraphPad Software). Tissue weights, food consumption and energy expenditure were analyzed by ANCOVA using XLSTAT (Addinsoft), with total body weight or lean mass as covariant.

## Author Contributions

Conceptualization, RKS, AK; Methodology, SV, JJ, PV; Formal Analysis, AK, IZ, SV, IHD, PRT, DAB, DH, RK, SF, PV, AS-P, JJ, RKS; Investigation, AK, IZ, SV, IHD, PRT, DAB, DH, RK, PV, AS-P, JJ, RKS; Writing – Original Draft, RKS, AK, IHD; Writing – Review & Editing, IZ, SV, PRT, DAB, DH, RK, AVP, SF, PV, AS-P, JJ, SO; Supervision, RKS; Project Administration, RKS; Funding Acquisition, RKS, SO, AVP

## Acknowledgements

RKS and SO were supported by the Wellcome Trust [grants WT098498 and WT095515 respectively], and by the Medical Research Council (MRC) [grant MC_UU_12012/5]. AVP and SV were funded by the British Heart Foundation [grant RG/12/13/29853] and the MRC [MC_UU_12012/2]. Animal work was conducted in the MRC Disease Model Core [MC_UU_12012/5]. We are grateful for technical assistance from Dr. Amy Warner at the MRC Disease Model Core, Mr Keith Burling at the MRC MDU Mouse Biochemistry Laboratory, Mr Warner and Ms Phillips at the Histology Core and Drs Brian Lam and Marcella Ma at the Genomics and Transcriptomics core. All of them are funded by MRC under MRC_MC_UU_12012/5. We would also like to thank Mr Gregory Strachan at the Imaging Core, which is funded by the Wellcome Trust [Grant 100574/Z/12/Z] for assistance.

